# Treating the Network: Targeted inhibition of two specific microRNAs in the brainstem prevents the development of hypertension

**DOI:** 10.1101/2020.03.11.987966

**Authors:** Jonathan Gorky, Danielle DeCicco, Sirisha Achanta, James Schwaber, Rajanikanth Vadigepalli

## Abstract

We here test the concept that disease states may result not from a single cause but from small changes in a network that are collectively significant. We recently showed that development of hypertension (HTN) in the spontaneously hypertensive rat (SHR) model of human essential hypertension is accompanied by changes in microRNA expression levels in the brainstem tracking the development of HTN ^1,2^. This led to the hypothesis that preventing the change in microRNA levels could prevent the development of HTN. We propose that hypertension emerges from a network that has been pushed out of a normotensive equilibrium into a compensatory, pathological state. We show that small perturbations in the gene regulatory networks in the brainstem by selectively blocking two microRNAs highlighted in our previous results, miR-135a and miR-376a, is sufficient to prevent development of hypertension in the SHR model. This effect appears driven by only modest changes in the expression of rate-limiting genes, many of which are targets of these miRNAs, suggesting that the combination of genes that are targeted in the network is responsible for the effect. The demonstration that hypertension is an emergent property of an underlying regulatory network suggests that a new treatment paradigm altogether is needed.

**One Sentence Summary:** A brief summary of the main result of your paper, without excessive jargon.

## Introduction

Essential hypertension has a neurogenic component, which is now established to be characterized by a specific state of inflammation, and dysregulated angiotensin signaling ^3,4^. Brainstem regions including the nucleus of the solitary tract (NTS), or the principle integrative center for blood pressure control, the rostral ventrolateral medulla (RVLM), caudal ventrolateral medulla and others have shown to be affected ^5–10^. The NTS is essential in blood pressure set point control. Blood pressure has previously been shown to be renormalized upon several interventions into NTS, including antagonizing the angiotensin II (Ang II) signaling pathway, and blocking the leukotriene B4 receptor (Ltb4r) ^11–15^. The nucleus of solitary tract (NTS) integrates visceral sensory information with higher order projections from hypothalamic and limbic brain regions in order to coordinate the sympathetic and parasympathetic responses of the autonomic nervous system. From a process control perspective, the NTS controls the blood pressure set point and functions as a comparator of afferent information that modulates autonomic effectors controlling blood pressure ^10^. In neurogenic hypertension (HTN), the NTS is considered an essential control point that drives the development of pathology, especially in the spontaneously hypertensive rat (SHR) model ^2^.

Recently, a global microRNA study was performed to evaluate the microRNA expression landscape in both NTS and RVLM in SHR and age-matched WKY controls ^1^. The study provided insight into a small number of microRNAs that might act as gene-regulatory network modulators influencing both the Ang II and Ltb4r signaling pathways along with several other potentially crucial processes. In NTS, both microRNAs were expressed at higher levels during the development of hypertension in SHR corresponding to 10-12 weeks of age, when physiological increases in blood pressure are first observed ^1^.

Each of these microRNAs have been previously characterized in other disease models, but neither have been implicated in hypertension as potential therapeutics. microRNA-135a has been shown to be expressed in the brain, and to be enriched in astrocytes compared to other CNS cell types including neurons, microglia and endothelial cells ^16^. Classically, microRNA-135a has been implicated in several cancers including colorectal, prostate, melanoma and glioma ^17–23^. microRNA-135a was implicated in the leukotriene B4 pathway via its regulation of the 5-lipoxygenase activating protein (FLAP) in hypoxia ^24^, and miR-135a was anti-correlated to its predicted targets in a regulatory network that impacts both the Ang II pathway and the Ltb4r pathway ^1^. miR-376a has also been shown to be expressed in the brain, and particularly enriched in neurons compared to the other cell types ^16^. microRNA-376a was implicated in hypertension through its predicted modulation of the Ang II pathway ^1^. Aside from its implication in neurogenic hypertension, microRNA-376a has been shown to be involved in apoptosis and DNA damage pathways ^25,26^, and it was also been characterized as a biomarker for type 2 diabetes and obesity ^27^.

Since both of these microRNAs displayed increased expression levels in NTS at the hypertension onset stage, corresponding to the 10-12 week age range, antagonizing these microRNAs at this critical time period was predicted to reduce the blood pressure in SHR via decreasing the activity of the Ang II and Ltb4r signaling pathways ^1,28^. The time period where these microRNAs display expression differences corresponds to a critical time period previously identified where intervention with losartan, a partial Ang II receptor antagonist, acutely attenuated hypertension in rats for up to a year ^29^. Moreover, microRNAs have been delivered intranasally in pre-clinical studies to act in the central nervous system, increasing their therapeutic value as CNS drugs ^17^. Therefore, the primary aim of the current study was to test whether transient inhibition of miR-135 and mir-376a combined, or inhibition of miR-135a and miR-376a alone would reduce blood pressure and restore molecular signaling network balances in SHR with minimal blood pressure and molecular effects in WKY.

## Results

### Single cocktail injection of anti-miR-135a and anti-miR-376a reduces blood pressure in SHR

Blood pressure in the SHR is increases between 10-12 weeks of age, yet a single ICV injection of a cocktail of anti-miR-135a and anti-miR-376a locked nucleic acids (LNA) at 10 weeks of age led to significantly decreased blood pressure over this same time frame (Figure 1, N=4-6 for each group). One week after LNA cocktail injection, the mean arterial pressure (MAP) decreased from an average of 165±2.5 mmHg to an average of 103±15.7 mmHg (mean±SEM), a 37.5% decrease from baseline (p<0.001). Two weeks after injection of the LNA cocktail, the MAP remained significantly decreased, with an average of 117±5.6 mmHg, a 29.1% decrease from baseline (p<0.01). Injection of a scrambled LNA oilgo not targeted towards any known miRNA or mRNA transcripts (miRCURY LNA inhibitor control, Exiqon) showed no significant changes from baseline nor from no surgery control SHR animals from the same cohort (Figure S2). The effects of individual LNA inhibitors were also tested, each having a smaller yet significant effect on lowering MAP in SHR (Figure S3).

**Figure 1:**
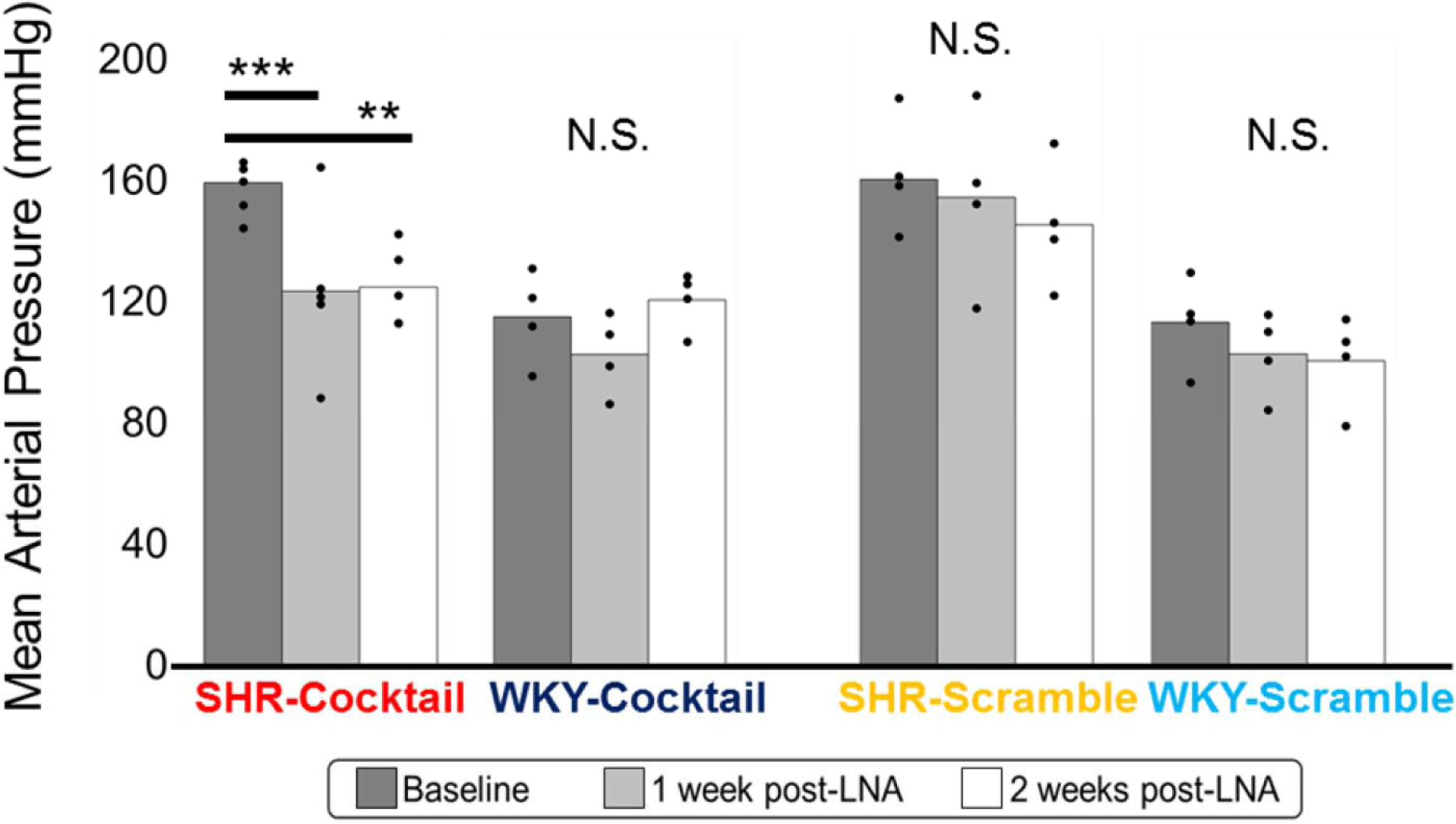
A single intracerebroventricular injection of a cocktail of anti-miR-135a and anti-miR-376a LNA renormalized blood pressure in the SHR for up to two weeks, but not WKY. N=4-6 for each group, ** p<0.01, *** p<0.001 (see supplemental methods for description of blood pressure acquisition and statistical calculations).

### Cocktail injection of anti-miR-135a and anit-miR-376a does not affect blood pressure of WKY

In contrast to the blood pressure altering effects of the anti-miR135a and anti-miR376a LNA cocktail on SHR, there were no significant effects on the blood pressure of WKY animals that serve as normotensive controls from a similar genetic background (Figure 1). Age-matched WKY rats that received a single central injection of the same cocktail of anti-miRs showed stable MAP averaging 99.0±8.9 mmHg at baseline and 99.5±7.8mmHg one week post injection (Figure 2, n=3-4 for each group). The scrambled LNA also did not show an appreciable effect on MAP, averaging 98.4±8.2mmHg at baseline and 101.5±5.7 mmHg (mean±SEM). You may include page breaks if you would like to embed the figures within the text instead of at the end of the paper.

**Figure 2:**
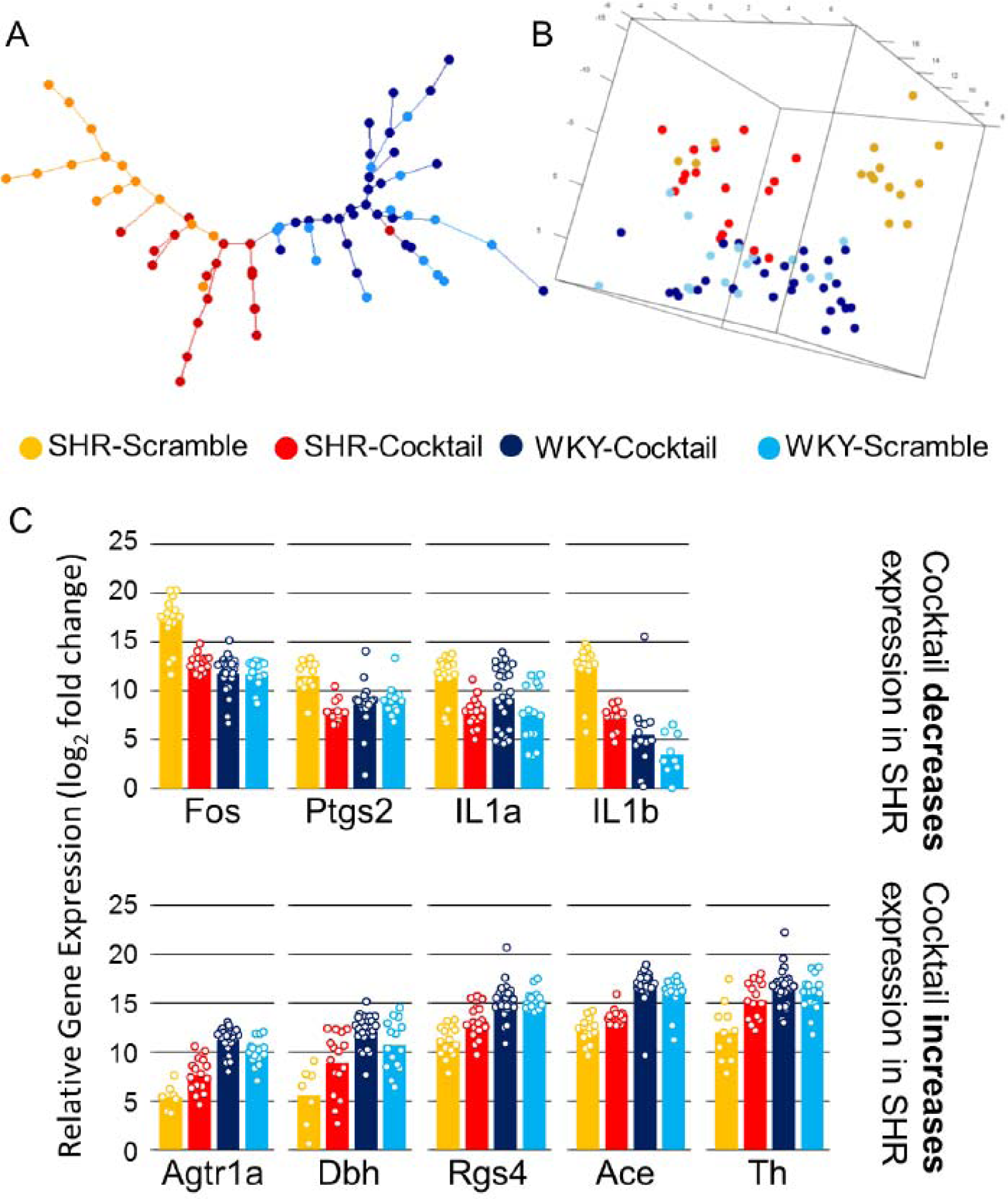
Injection of the LNA cocktail causes NTS gene expression in the SHR to become more similar to that of WKY. Minimum spanning tree (A) principal component analysis plot (B) and showing the similarity of individual NTS samples taken across each cohort of animals. In both cases, the SHR-cocktail cohort has greater similarity to WKY than that of the SHR-scramble cohort, suggesting a molecular substrate for the observed physiological changes in SHR-cocktail cohort. Some of the genes showed decreased expression in response to LNA cocktail injection and others showed increased expression (C). Statistics for all comparisons are given in Figure S5. Gene expression data for all other genes measured including reference genes can be found in Figure S4.

### Anti-microRNA cocktail nudges SHR gene regulatory network closer to that of WKY

Using a cohort of 19 genes related to inflammatory processes, angiotensin signaling, and autonomic function, we found that the perturbations in the gene regulatory networks of the nucleus tractus solitarius (NTS) were indeed prevented by injection of the LNA cocktail, but only in SHR and not in WKY. A minimum spanning tree (Figure 2A) based on the expression values of this cohort of 19 genes shows individual laser captured NTS samples taken across all animals in the LNA injection study. The effect of the anti-miR LNA cocktail on SHR shows a shift toward the WKY gene expression pattern, without fully recapitulating it. The effect of the anti-miR LNA cocktail on WKY is not distinguishable from the negative control scramble LNA, which mirrors the observed physiology. To test that this result was not an artifact of the data analysis method, we also performed a principal component analysis (PCA) using the same dataset. The PCA plot (Figure 2B) shows a similar trend of separation with the SHR-scramble cohort generally clustered away from the SHR-cocktail and both WKY cohorts. In the principal component space, the SHR-cocktail group more closely resembles that of both WKY cohorts than it does the SHR-scramble cohort.

The gene expression profiles also reflected a general trend of the anti-miR LNA cocktail shifting the SHR expression pattern to be more in line with that of WKY (Figure 2C). While several bioinformatics-predicted microRNA targets showed expression changes in the SHR in response to anti-miR LNA cocktail injection (Figure 2C), a few predicted target genes did not show differential expression response to the LNA cocktail (Figure S4). While some of the gene expression changes shown do not meet the traditional, arbitrary criterion of p<0.05, the anti-miR induced adjustments in gene expression accumulate into an overall trend of a clear shift in SHR gene expression pattern towards that of WKY. The full table of p-values is given in Figure S5.

While Figure 2C shows genes whose expression values were affected by the cocktail injection in SHR, there were other genes measured whose values were not affected by the cocktail injection. Still none of the genes showed a significant difference between WKY-cocktail and WKY-scramble. Renin and Alox5ap had higher levels in SHR than WKY without a large change in expression a result of cocktail treatment. Agtrap showed statistically significant, yet very small changes in response to the cocktail treatment in SHR, but these levels are not distinguishable from the WKY average. The levels of Ptgr1 are the same across all SHR and WKY cohorts. IL10, Agtr1a, and Jun showed lower levels in the SHR cohorts than in the WKY cohorts and were unaffected by the cocktail treatment (Figure S4). The statistics for all comparisons of gene expression values two weeks after LNA injection are given in Figure S5. The p-values shown were determined by fitting the data to a mixed linear model ANOVA with Satterwaithe degree of freedom calculations to account for multiple measurements taken for each animal. This was done using the LMER package in R. In this table, p-values below the traditionally accepted cut-off of p<0.05 are bolded in this table, although this should not be construed to assume the authors consider this to be a viable cut-off for reproducible results. By considering the changes in aggregate is perhaps more useful for showing that the network is affected rather than one or another of its constituents.

The statistics for all comparisons of gene expression values two weeks after LNA injection are given in Figure S5. The p-values shown were determined by fitting the data to a mixed linear model ANOVA with Satterwaithe degree of freedom calculations to account for multiple measurements taken for each animal. This was done using the LMER package in R. In this table, p-values below the traditionally accepted cut-off of p<0.05 are bolded in this table, although this should not be construed to assume the authors consider this to be a viable cut-off for reproducible results. By considering the changes in aggregate is perhaps more useful for showing that the network is affected rather than one or another of its constituents.

### NTS of SHR and WKY have inherently distinct gene expression networks

In a direct examination of the relationship between mircoRNAs and gene networks in neurons of the NTS, we have obtained simultaneous expression values for a cohort of 52 genes along with miR-135a and miR-376a in laser captured pooled single neurons from SHR and WKY animals. This effort generated nearly 4,700 data points that were used to examine gene co-expression networks. These networks reveal a surprising difference between the relationships of miR-135a and miR-376a to the genes measured, most notably that the microRNA-gene expression correlation patterns between SHR and WKY are distinct and that both microRNAs are more strongly connected in the SHR network compared with WKY network (Figures 3, S6, S12). This suggests a basis for why there were SHR-specific effects of anti-miR LNA cocktail on gene expression and physiology; the homeostatic regulatory networks that these microRNAs operate in are different between the two strains.

**Figure 3:**
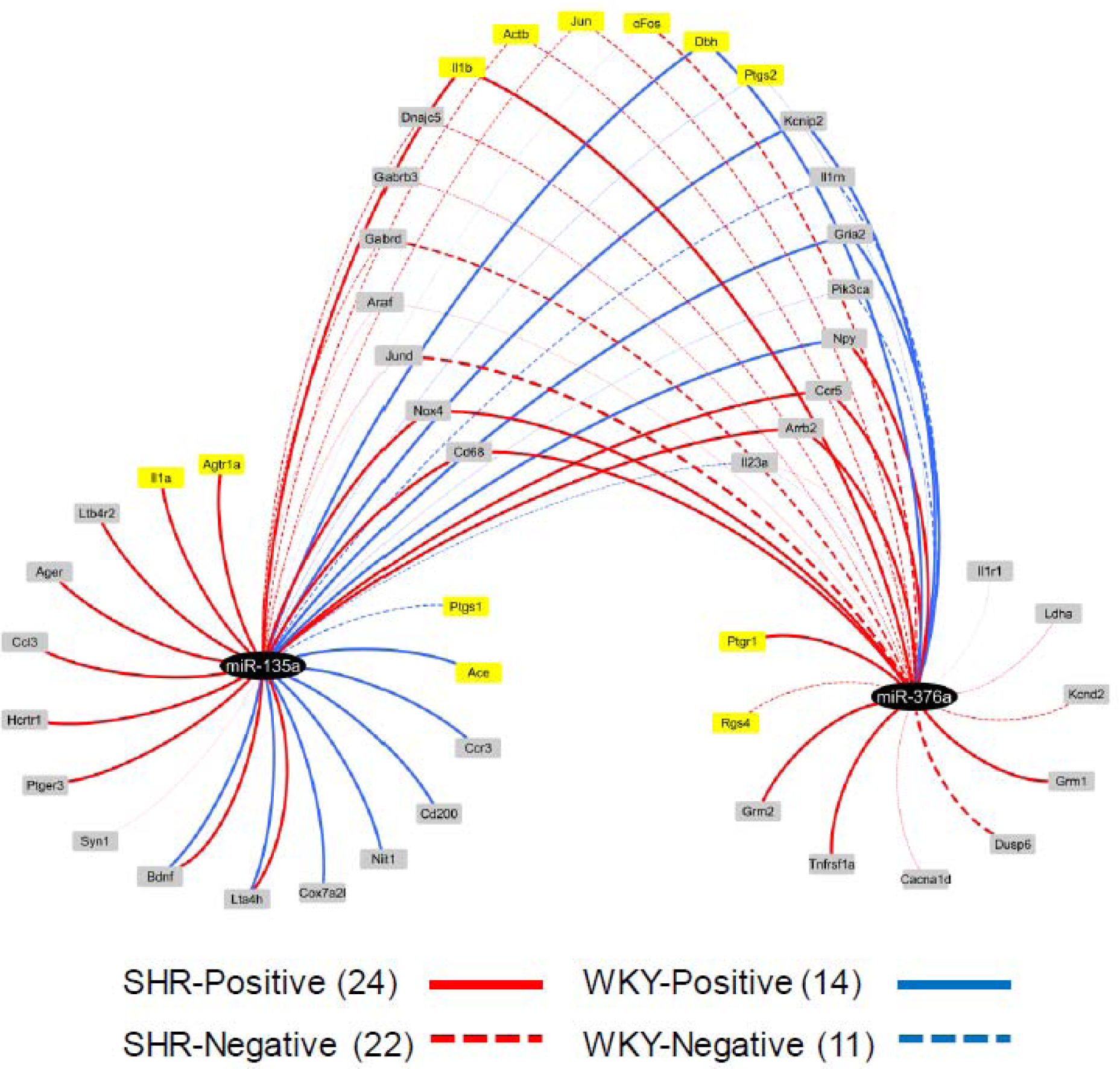
The co-expression networks for miR-135a and miR-376a are distinct in NTS neurons of SHR and WKY animals, with the connectivity in SHR being nearly twice that of WKY for genes related to inflammation, angiotensin signaling, and autonomic function. This suggests an underlying rationale for why LNA cocktail injection affected only SHR and not WKY both physiologically and at a transcriptional level. Gene co-expression network derived from laser captured NTS neurons across N=4 SHR and WKY animals. Edges represent Pearson correlation R values, only correlations that met a q value cut-off of q<0.05 are shown. Nodes in this figure are genes that demonstrated significant correlations with either or both miR-135a or miR-376a. Genes not found to be significantly correlated to these microRNAs were omitted from the figure. Nodes colored in yellow are genes that were also measured in the LNA treated animals two weeks after injection, as shown in Figure 2C and S4. The numbers in parenthesis in the legend are counts of the number of edges for each type of interaction.

## Discussion

When considering the network of organs that drive and perpetuate hypertension, the NTS is a node with crucial connectivity ^2^. Through its role in interpreting interoceptive inputs from the entire body and translating that into autonomic function via the parasympathetic and sympathetic outflows, it forms an attractive target for manipulation. Past studies have shown that manipulations affecting the NTS alone are sufficient to ameliorate hypertension in the SHR model ^3,30–33^. Many of the pharmacological interventions that show clinical efficacy in lowering blood pressure in hypertensive patients work at many different sites of action in parallel, even if the presumed mechanism of action is narrowly defined. Consider, as an example the angiotensin receptor blocker losartan, whose mechanism of action generally is understood as working on angiotensin-1 receptors in the heart, vasculature, and kidney, but which also has effects in the brain that were not initially appreciated. Such interventions also have the tendency to drive a very large direct effect on only one receptor or protein, which then has downstream consequences that lead to renormalizing blood pressure.

The use of microRNA inhibitors, however, did not lead to large, restricted changes in the network. Any given microRNA has tens to hundreds of putative binding sites on known mRNA transcripts and plays only a relatively small role in the overall expression levels of that transcript8. Rather than expecting microRNAs to be modulators of select gene expression, it would perhaps be more functionally useful to consider microRNAs as tuners of distributed gene regulatory networks. Through small-in-effect but broad-in-scope changes in gene networks, it is possible to retune the network behavior that leads to distinct physiology, as evidenced by the results shown in Figures 1 and 2. These results form the basis for follow-on studies focused on detailed unraveling of the molecular mechanisms that drive this significant physiological outcome of preventing hypertension. Considering the network-wide small changes observed in this study, we propose that a network-based approach that targets several components is essential to unraveling detailed mechanisms of how in vivo microRNA manipulations alter organismal physiology. In this regard, obtaining global transcriptional changes over time and evaluating these changes using computational models of the dynamic influences between regulators and targets is likely to yield fruitful follow-on directions ^2^. The initial results presented here suggest that knockdown of individual microRNAs may not necessarily lead to very large changes in the level of any individual transcript, but that the expression of a number of distinct transcripts is affected. This is in line with a recent theoretical proposition that that physiological function is the product of network interactions and that small changes in such networks can lead to drastically different physiological outcomes ^34^.

Instead of thinking about microRNA as modulators of gene expression, it may be more functionally useful to consider them to biological process modulators. The work presented here in concert with a growing literature demonstrating manipulation of microRNA is sufficient to have robust physiological effects makes this concept plausible. Especially given the relatively low effect size in gene expression or translation of any given microRNA interacting with its cohort of transcripts, it is challenging to image evolutionary pressures acting to generate such modulators unless there were biological process level effects, at least in some cases. The data presented here, where anatomically targeted manipulation of two microRNAs was enough to have broad effects on physiology.

Neurogenic hypertension is often refractory to current therapeutic paradigms that directly affect only one or a few targets at a time ^10^. We have shown that modulating microRNAs can renormalize a broad network through direct effects on expression of genes coding for proteins that are the rate-limiting steps in renin-angiotensin, autonomic, and inflammatory pathways. Further, when simultaneously targeting many nodes in that network, relatively small individual effect sizes on each node appears to be sufficient to drive the rest of network into a normotensive state. These results suggest a path forward in the development of therapeutics for disease states where single target interventions have failed. To treat a disease that emerges out of a complex network, we must find ways to treat the network.

## Materials and Methods

### Animal Model

Male, Wistar Kyoto (WKY/NHsd) rats and spontaneously hypertensive rats (SHR/NHsd) obtained from Harlan Laboratories (Now Envigo) were housed 1 per cage in the Thomas Jefferson animal facility to avoid animal to animal stress from dominance that could affect blood pressure. Facilities were maintained at constant temperature and humidity with 12/12 hour light/dark cycles. All protocols were approved by the Thomas Jefferson University Institutional Animal Care and Use Committee.

### Laser Capture Microdissection (LCM)

Laser capture microdissection was used to obtain precise regional samples from NTS. To preserve RNA integrity, dehydration was performed within 20 minutes. Slides were fixed in cold Acetone with 1% Hydrogen Peroxide (Sigma-Aldrich) for 1 minute, then rinsed in phosphate buffered saline (PBS) for 30 seconds. Slides were then dehydrated in graduated ethanol concentrations followed by two immersions in xylene for 1 minute and then 5 minutes. The LCM process was performed using Arcturus XT LCM attached to a Nikon TE2000 microscope. The NTS region coextensive with the area postrema was localized using anatomical landmarks, and was lifted as a region using the laser capture dissection method. Several (2-4) bilateral NTS sections were pooled per each cap, and sections were lysed directly on the cap, and cooled on ice before -80 degree Celsius storage.

### microRNA and mRNA simultaneous high-throughput real-time quantitative PCR

microRNA expression levels and transcript expression levels were measured using a sequential reverse transcription reaction from one LCM-obtained or RNA sample. Samples of bilateral NTS obtained from LCM and lysed on cap were directed into universal reverse transcription reaction to reverse transcribe all mRNA (VILO, ThermoFisher). Following a universal reverse transcription, the sample underwent reverse transcription for specific microRNAs (miR-135a and miR-376a) using the Taqman detection system to increase specificity of future quantification. Due to limited RNA amount, one universal preamplification step was performed using primers for genes of interest and microRNAs of interest using TaqMan® PreAmp Master Mix per the manufacturer’s protocol (Applied Biosystems, Foster City, CA). Finally, genomic DNA was digested and genes and microRNAs were measured using multiplex RT-qPCR (BioMark by Fluidigm) or standard EvaGreen-based qPCR (ABI 7000).

### Multiplex RT-qPCR

For the LNA treated animals, each sample was bilateral punch of the NTS from a 200um brainstem slice. Animals were sacrificed via rapid decapitation without anesthesia and brains removed and sliced at 200um and kept on ice cold aCSF until punches were taken with the aid of a surgical microscope. Samples were then put into lysis buffer; the time from sacrifice to placing the sample into lysis buffer took no more than five minutes. Samples were processed using the RNeasy Mini-Prep (Qiagen) kit according to the manufacturer’s recommended protocol. After extraction, total RNA was quantified using the NanoDropTM system. Each sample’s RNA was reverse transcribed to cDNA using SuperScript III VILO Master Mix (ThermoFisher) according to manufacturers recommended protocols.

For the transcript measurement and quantification for the previously prepared sample, qPCR reactions were performed using a combination of 96.96 and 192.24 BioMark™ Dynamic IFC Arrays (Fluidigm®, South San Francisco, CA) enabling quantitative measurement of multiple mRNAs and samples under identical reaction conditions (Spurgeon et al. 2008). Each run consisted of 30 amplification cycles (15s at 95°C, 5s at 70°C, 60s at 60°C). Ct values were calculated by the Real-Time PCR Analysis Software (Fluidigm). Intron-spanning PCR primers for every assay were designed (Figure S15) and tested for proper targeted amplification before use on a BioMark array.

### Data Normalization for Multiplex RT-qPCR

We followed the methods we have previously utilized for high-throughput transcript level analysis^1^. In brief, assays and samples failed by software due to technical noise were discounted for downstream analysis, and data was normalized by median gene expression levels across 24 genes in the case of 192.24 dynamic array and across 96 genes in the case of the 96.96 dynamic array. Several reference genes were also checked for their ability to normalize data against and were found to highly correlate to the sample median across all viable gene assays.

### Blood pressure measurements

Blood pressure measurements were done with the noninvasive CODA VPR tail cuff blood pressure measurement system (Kent Scientific, Inc). Rats were acclimatized to the restraint devices and the blood pressure measurement procedure for several weeks prior to taking baseline and experimental readings. Rats were gently coaxed into the restraint devices and placed in a heating pad for five minutes prior to any measurements. The ambient temperature was maintained at 79 – 84 °F in order to promote blood flow through the tail without causing stress due to overheating. The BP measurement procedure has 5 cycles for acclimation followed by 20-30 cycles of actual measurements. For each cycle, the tail blood flow volume, heart rate, systolic, and diastolic blood pressure are measured. Cycles with a tail flow volume greater than 5 mL/min were considered to have registered the VPR sufficiently to obtain a reading. The set of cycles from each session that was used was determined by taking the longest consecutive run of “good” measurements. A mixed linear model ANOVA was used to analyze the blood pressure measurements for each time point in order to find significant differences between treatment groups over time (lme4 package in R).

The volume pressure method of rodent blood pressure measurement has been shown to strongly correlate with blood pressure measured by telemetry. The VPR system does tend to over-estimate very low blood pressure and under-estimate very high blood pressure. However, given these tendencies, the expectation would be that the dynamic range of the VPR would be compressed and would thus be more likely to not find significant differences if they existed.

### Stereotaxic Injections of Locked Nucleic Acids

The 25 uL injection consisted of a 1:1 ratio of anti-miR-376a and anti-miR-135a Locked Nucleic Acid (LNA, Exiqon) 10 uM solution in artificial cerebrospinal fluid, this condition was considered AM-Combo. A negative control antisense oligonucleotide LNA (a sequence that does not align to any known microRNA or mRNA transcript), called “Scrambled”. No surgery and sham surgery conditions were also assayed. Sham surgeries included cannulas that were inserted and an empty needle was placed for the duration of the corresponding injection time were also performed. The rats were anesthetized by isoflurane inhalation and placed in a stereotactic instrument (Stoelting Co., Wood Dale, IL, USA), positioned, and the incision site appropriately prepared. A small incision was made, and the skull was then exposed and cleaned and a small burr hole was drilled using a dental drill or Dremel rotary tool, and cannula was mounted with dental cement to skull screws. The incision was closed and the rats were placed on an isothermal pad at 37°C and continuously observed following surgery until recovery. Topical analgesic was applied immediately following the surgical procedure and as needed thereafter. The LNAs were injected using a Hamilton syringe with a 35-gauge needle (Reno, NV, USA) over a 30 minute time period into the fourth ventricle using the coordinates: 11.3 mm caudal to bregma, 0.5 mm lateral to midline, and 7 mm below the surface of the skull.

Six SHR were injected with AM-Combo, three were injected with AM-376a, three were injected with AM-135, and four SHR were injected with scrambled oligonucleotide control. Two SHR underwent sham surgery where a cannula was placed, but no injection was given. Four SHR were maintained with the cohort as untreated. Four WKY were injected with AM-Combo and four were injected with scrambled negative control.

At the end of each time course of treatment, rats were sacrificed via rapid decapitation, and brainstems were excised and placed into Optimal Cutting Medium (OCT, TissueTek, QIAGEN, Valencia, CA) and flash-frozen at -80 degrees Celsius for cryosectioning. Brainstem samples were taken throughout the intermediate levels of the NTS co-extensive with the area postrema via laser capture microdissection.

### Statistical and Analytical Methods for in vivo gene expression after LNA injection

After quality control and normalization of the multiplex RT-qPCR data, significant differences in expression values from NTS samples taken across all animals were fit to a mixed linear model with Satterwaithe degree of freedom calculations to account for the differing number of multiple measurements per animal. Several genes showed significantly different expression values as given in the supplemental table B. Such annotation was left out of the bar plot figure in order to minimize visual crowding. A consideration of the aggregate differences between SHR – Cocktail, SHR – Scramble, and WKY – Average using analysis of principal components was able to demonstrate the differences between SHR – Scramble and the other two groups across all genes aggregated were significant at the p<0.05 level. Differences between SHR – Cocktail and WKY – Average were not significant.

Bar plots from Figure 3 of the main paper for the gene expression data show the average values across all animals with error bars showing the standard error of the mean, determined at the animal level using error propagation to account for multiple measurements. The spanning tree in Figure 2 was generated using the vegan package in R. The distance matrix for spanning tree was generated using Euclidean distance across all genes measured. The principal component plot in Figure 2 was generated using the pcaMethods package in R with the ‘rnipals’ method for interpolation.

**Supplemental Figure S1:**
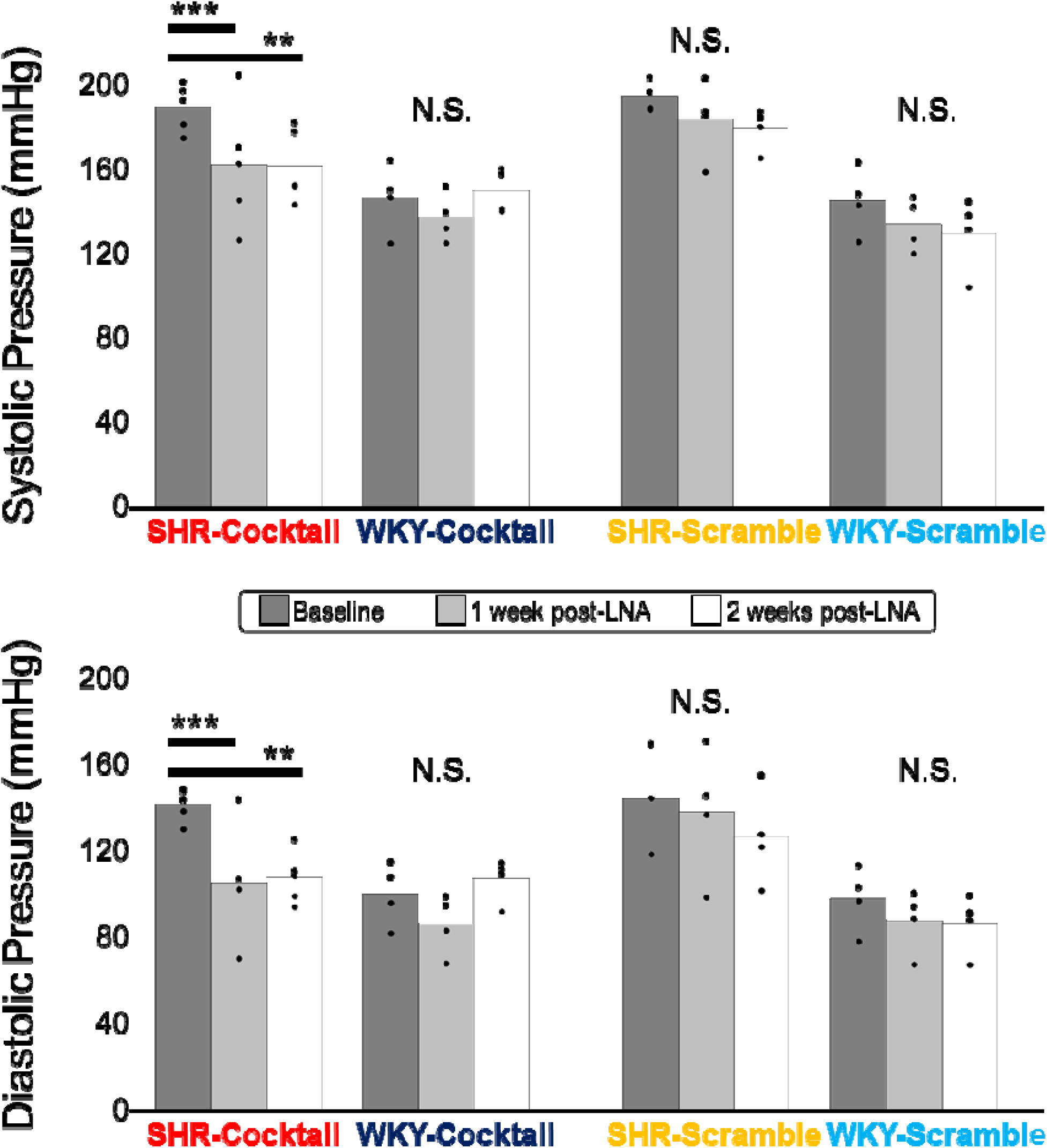
Systolic and diastolic blood pressure are both lowered from anti-microRNA LNA injections. This figure shows the corresponding components of blood pressure from the same experiments shown in Figure 1 of the main paper. N=4-6 for each group, * p<0.05, ** p<0.01, *** p<0.001 (see supplemental methods for description of blood pressure acquisition and statistical calculations). Error bars represent standard error of the mean.

**Figure S2:**
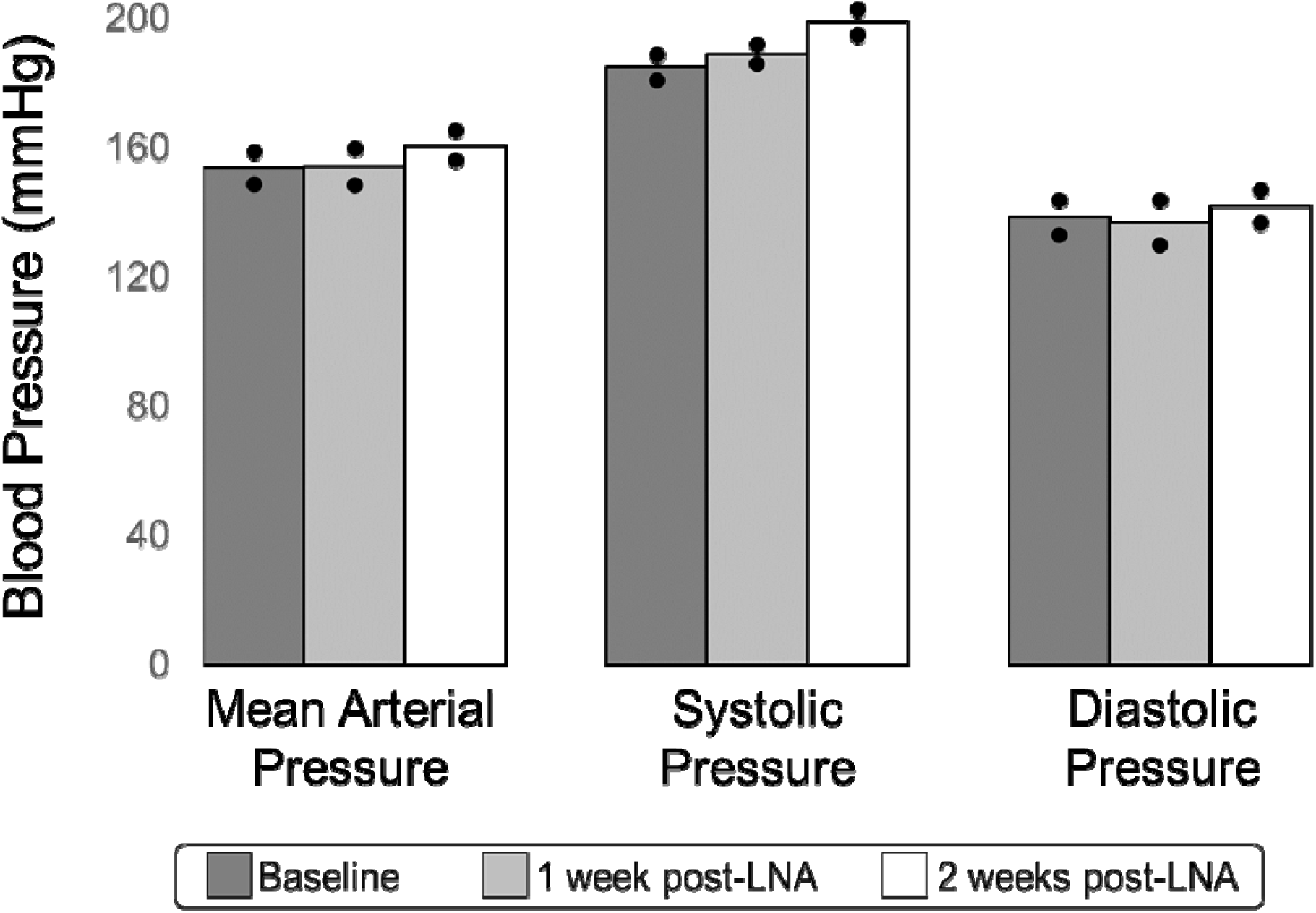
Data from SHR animals in a sham surgical condition, where a cannula was placed, but no subsequent injection was made. The sham surgery SHR showed no significant changes (N=2, nested ANOVA with Tukey post-hoc to account for multiple measurements) from baseline as a result of having the cannula placed into the 4^th^ ventricle without injection.

**Figure S3:**
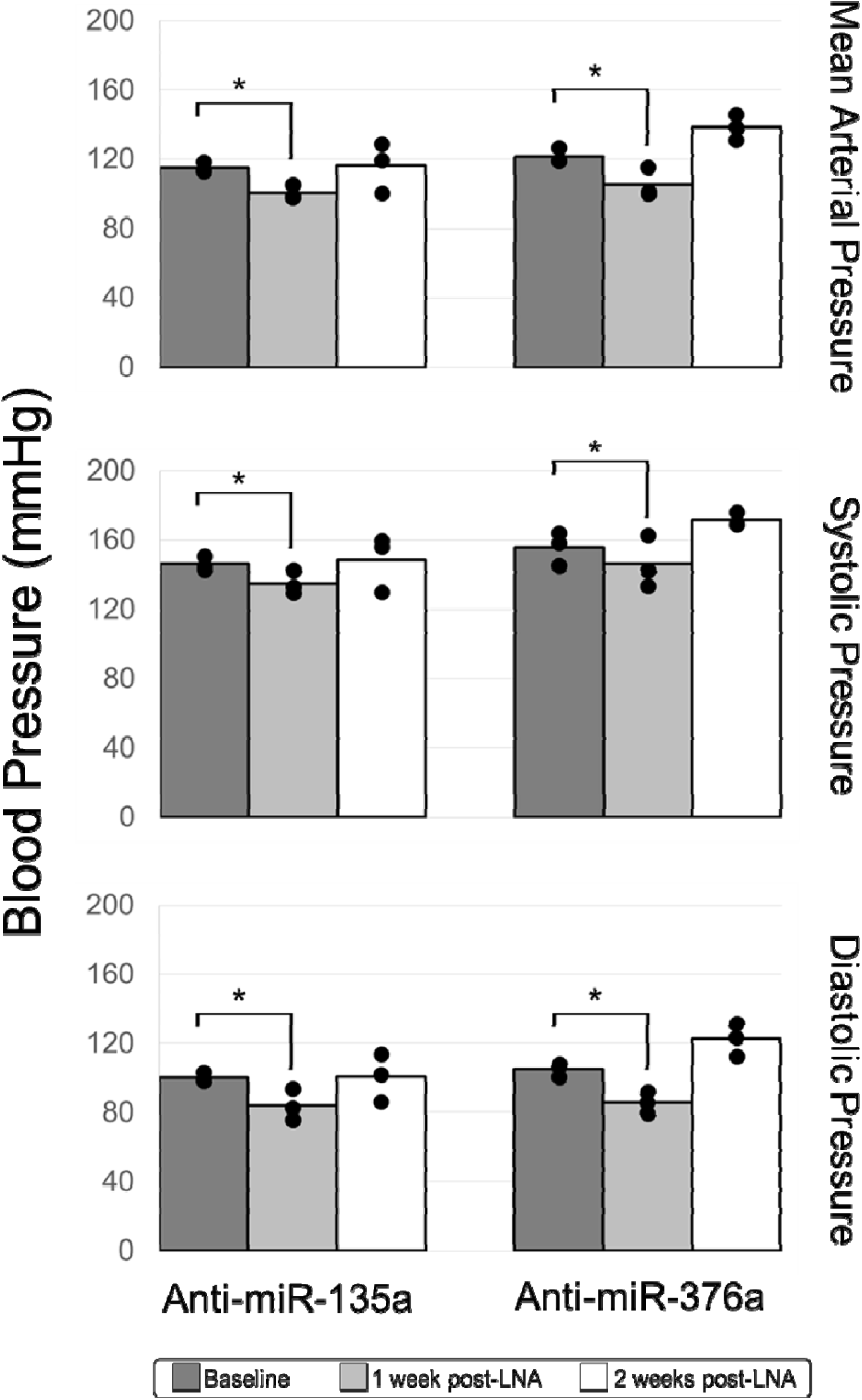
Injection of either anti-miR-135a or anti-miR-376a caused a significant decrease blood pressure, but with an effect size much smaller than the two given at the same time, as we done in the cocktail injections in Figures 1 and S1. N=3 for each group, * p<0.05 nested ANOVA with Tukey post-hoc.

**Figure S4:**
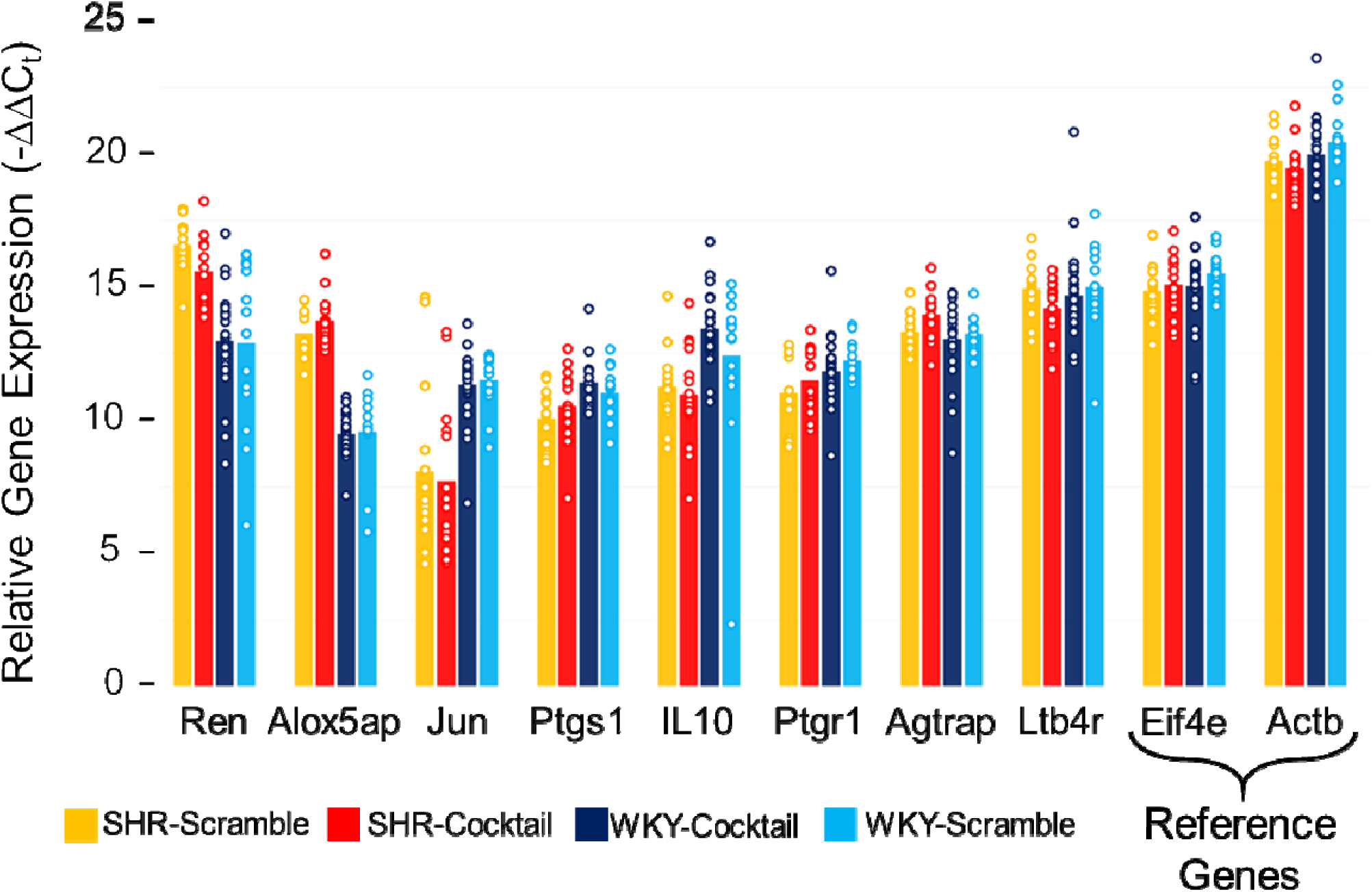
Several genes measured in the SHR and WKY cocktail injection experiments did not follow the pattern of renormalization that as seen in Figure 2C. The renormalization pattern is one where the SHR-Cocktail values have moved away from the SHR-Scramble and closer to the WKY-Combined. Statistics for all group comparisons of genes shown here and from Figure 2C can be found in Figure S5.

**Figure S5:**
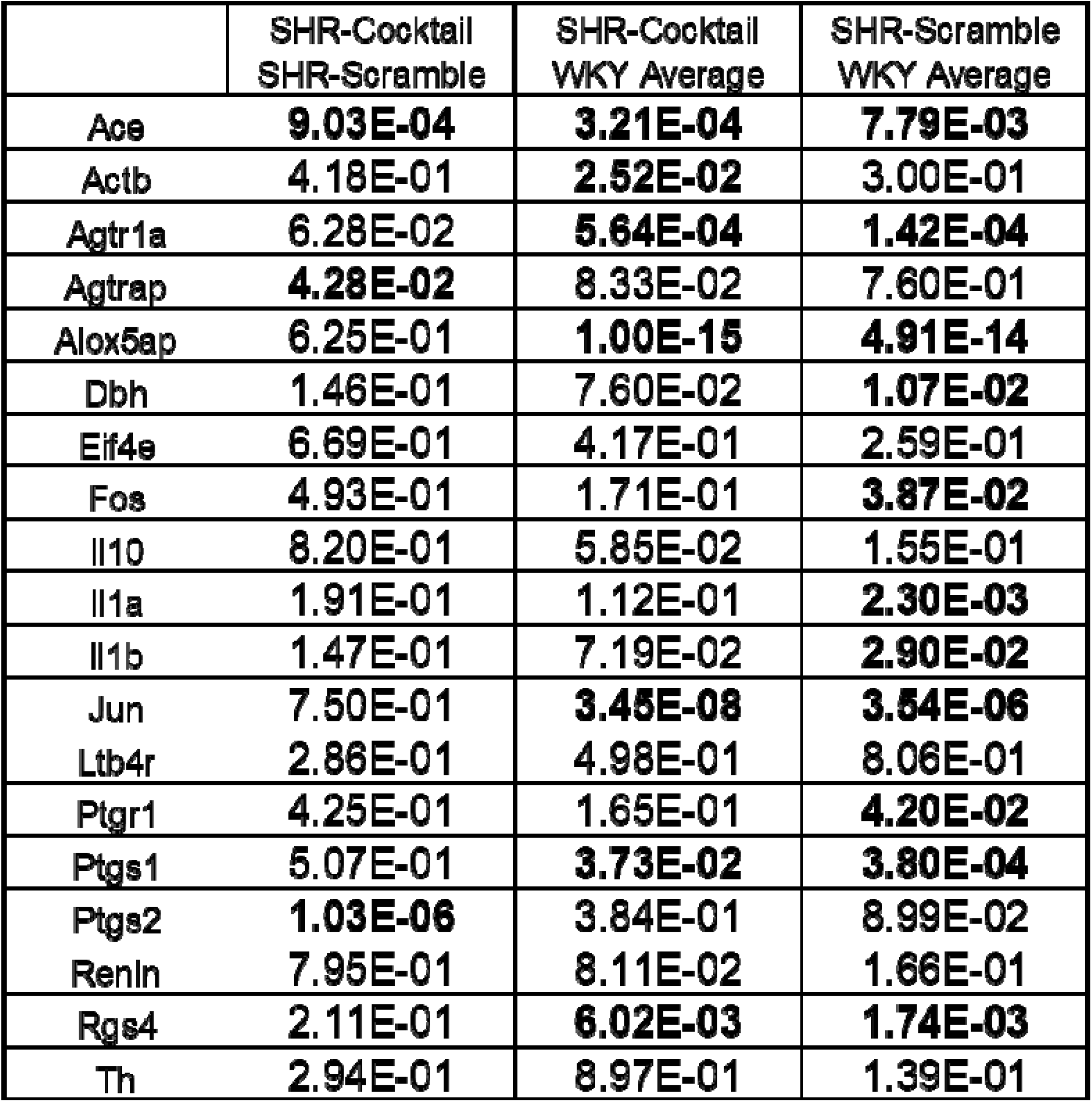
Table of p-values from comparisons of gene expression from all experimental groups two weeks after cocktail or scramble LNA injection. These p-values were computed using a mixed linear model ANOVA with Satterwaithe degrees of freedom (LMER package in R). Values below the traditionally accepted significance cut-off of p<0.05 are bolded in the table.

